# Kinesin-8/Kip3 requires beta tubulin tail for depolymerase activity

**DOI:** 10.1101/2025.01.27.635192

**Authors:** Kaitlin Alemany, Samantha Johnson, Jeffrey K. Moore

**Author notes:** Contact information for corresponding author: Jeff Moore, MS8108, 12801 E 17th Ave, Aurora, CO 80045.

## Abstract

Carboxy-terminal tails (CTTs) of tubulin proteins are sites of regulating microtubule function. We previously conducted a genetic interaction screen and identified Kip3, a kinesin-8 motor, as potentially requiring the β-tubulin CTT (β-CTT) for function. Here we use budding yeast to define how β-CTT promotes Kip3 function and the features of β-CTT that are important for this mechanism. We find that β-CTT is necessary for Kip3 depolymerase activity, but not for microtubule binding and motility. Mutant yeast cells lacking β-CTT show increased accumulation of Kip3 at plus ends and along microtubules, but no increase in catastrophe when Kip3 is overexpressed. *In vitro* experiments show that the β-CTT is necessary for Kip3 to form a tight complex with soluble tubulin but is unnecessary for Kip3 to bind tubulin in the microtubule lattice. These results suggest a model in which β-CTT promotes Kip3 depolymerase activity by supporting a Kip3-tubulin binding state that is only accessible at the microtubule plus end or in solution.

## Introduction

Cells create complex and diverse microtubule (MT) networks through combinations of extrinsic and intrinsic regulation. MTs are polarized, cylindrical polymers composed of α/β-tubulin heterodimers that exhibit dynamic instability, where the end of the polymer (typically the plus end) transitions between states of polymerization and depolymerization^1–3^. This behavior is controlled by the ability of tubulin to visit straight (MT-favoring) and curved (MT-disfavoring) conformations. Microtubule associated proteins (MAPs) act as extrinsic regulators^1,4^. For example, different kinesin motors directly regulate MT dynamics via binding and altering tubulin conformation, or indirectly via transporting other MAPs to MT ends^1,4,5^. The α- and β-tubulin subunits act as intrinsic regulators. α- and β-tubulins are encoded by multi-gene families known as isotypes, which are differentially expressed according to metazoan cell type and diverge across species. While the globular body of tubulin proteins have high sequence conservation across isotypes and often species, the disordered and acidic carboxy-terminal tail (CTT) domains that decorate the MT surface are highly divergent^6,7^. Tubulin CTTs are major sites for MT-MAP interactions and for post-translational modifications (PTM) providing an intersection of intrinsic and extrinsic regulation that is thought to create tunable programs for cargo transport and MT dynamics. This model is referred to as the ‘tubulin code’^1,6,8^. Analogous to how the ‘histone code’ regulates the architecture and interactions on chromatin^9^, the tubulin code regulates the interactions and activities of various MAPs on the MT surface^1,10,11^. While the field has a strong understanding of the tubulin genes and modifications that form the basis for the tubulin code, we have a comparatively weak understanding of how CTTs interact with MAPs that ‘read’ the code and how differences in CTTs give rise to network-level complexity.

Previous studies have identified several motor proteins that are sensitive to changes in CTTs^s4,11,12^. Experiments with purified yeast tubulin showed that both the α and β-tubulin CTTs promote the velocity and processivity of kinesin-1. This same study showed that only β-CTT promotes dynein processivity, but has no effect on velocity^4^. These results show that the α- and β-CTTs cause distinct effects on different motors. A study using purified mammalian tubulin to study the CTT PTM, polyglutamylation, showed that kinesin-1 run length was decreased by β-CTT glutamylation and was insensitive to modifications on the α-CTT^12^. Further studies of CTT function *in vivo* in budding yeast showed that two motors from the kinesin-5 family, Cin8 and Kip1, are differentially regulated by the β-CTT with β-CTT promoting plus-end motility of Cin8^11^. Collectively, these findings demonstrate that varying the composition of tubulin CTTs can create effects on motor function that may be specific to subsets of kinesin or dynein motors. However, only a small number of kinesin motors have been tested for CTT sensitivity, and it remains unclear how CTTs differentiate between motors for their selective effects. Which aspects of the kinesin activities are altered by the CTTs? Is the mechanism the same across classes of kinesins or are there class-specific features that enable some motors to differentially ‘read’ CTTs?

To begin investigating the mechanism for how motors differentially read CTTs, our lab previously conducted a genetic screen to identify roles for CTTs in budding yeast^10^. The results indicated a loss of Kip3 function in mutants that lack the β-CTT. Kip3, a kinesin-8 family motor, regulates MT dynamics by combining its two main activities: processive motility to MT plus ends and plus end depolymerase activity^5,13,14^. In the antenna model proposed by the Howard lab, as the MT grows to longer lengths, more Kip3 lands on the MT and walk towards the plus end, where the motors accumulate and promote depolymerization; accordingly, longer MTs recruit more Kip3 and are selectively targeted for depolymerization^5,13,15^. Recent structural studies suggest that Kip3 exhibits distinct conformations to bind to straight tubulin in the lattice versus the curved tubulin at the plus end, which is necessary for Kip3 induced depolymerization^16–18^. It is hypothesized that in the latter conformation, Kip3 locks the protofilament in a bent state that destabilizes the plus end, allowing Kip3-tubulin complexes to detach from the MT^16,18^. In this way, Kip3 acts as a plus-end depolymerase that promotes catastrophe^5,19^. Despite these studies, the mechanism behind how Kip3 can transit between states of binding to tubulin in the lattice, walking to the plus end, and switching to bind to curved tubulin for depolymerization remains unclear.

In this study we utilized the budding yeast *Saccharomyces cerevisiae* to investigate the relevance of tubulin CTTs for the Kip3 mechanism. Budding yeast express only two α-tubulin and one β-tubulin isotypes and lack PTMs, thus simplifying the components of the tubulin code^10^. We studied which function of Kip3 is affected by β-CTT with live-cell imaging in budding yeast where we can genetically manipulate CTT composition and visualize individual MTs emanating from spindle pole bodies (SPBs; the centrosome equivalent in yeast), and complementary *in vitro* reconstitution assays where Kip3 motors bind and walk along MTs made of native yeast tubulins. We find that β-CTT is required for Kip3’s depolymerase activity and binding to soluble tubulin, but not necessary for motility or binding to tubulin in the MT lattice. We propose that β-CTT could act as a tunable control for kinesin-8 activity across species or across cell types in metazoans. Our discoveries are consistent with the tubulin code hypothesis but suggest an additional dimension – CTTs may act differently at MT ends compared to MT lattices.

## Results

### Kip3 requires β-CTT for function

We previously conducted a genetic screen in yeast strains that lacked either α-CTT (*tub1*-442) or β-CTT (*tub*2-430Δ), predicting that CTT mutants should exhibit genetic interaction profiles that are similar to those of null mutations in MAPs that require CTTs for function^10^. The screen found 67 negative interactors for α-CTT and 247 for β-CTT. We used GeneMANIA to calculate a composite functional association network amongst the negative genetic interactions to predict co-functionality with additional genes outside of negative interactions^20^. Specifically, this prediction computes the degree of association these genes have with the list of negative interactions, sorted by biological, molecular, and cellular component Gene Ontology terms. Through this process, we identified *KIP*3, a kinesin-8 family member, as one of the top 10 genes likely to be co-functional with β-CTT. Kip3 was not identified from a similar analysis of α-CTT genetic interactors (data not shown). A previous genetic interaction screen conducted with a *kip*3Δ null mutant identified strong negative interactions with 84 genes (see Methods)^21^. 22 of these 84 genes are shared amongst the negative genetic interactions for the *tub*2-430Δ screen (Figure 1A). Sorting the shared genes by Gene Ontology revealed involvement in processes such as cytoskeletal motor activity, nuclear migration and establishment of mitotic spindle orientation (Figure 1B, S1A and B). Since the 22 genes become essential when either Kip3 or β-CTT are ablated, then it is likely that Kip3 and β-CTT are also necessary for these processes to occur. Collectively, these data suggest that Kip3 and β-CTT function in a common pathway(s), and that this pathway contributes to positioning the mitotic spindle and nucleus during cell division.

**Figure 1:**
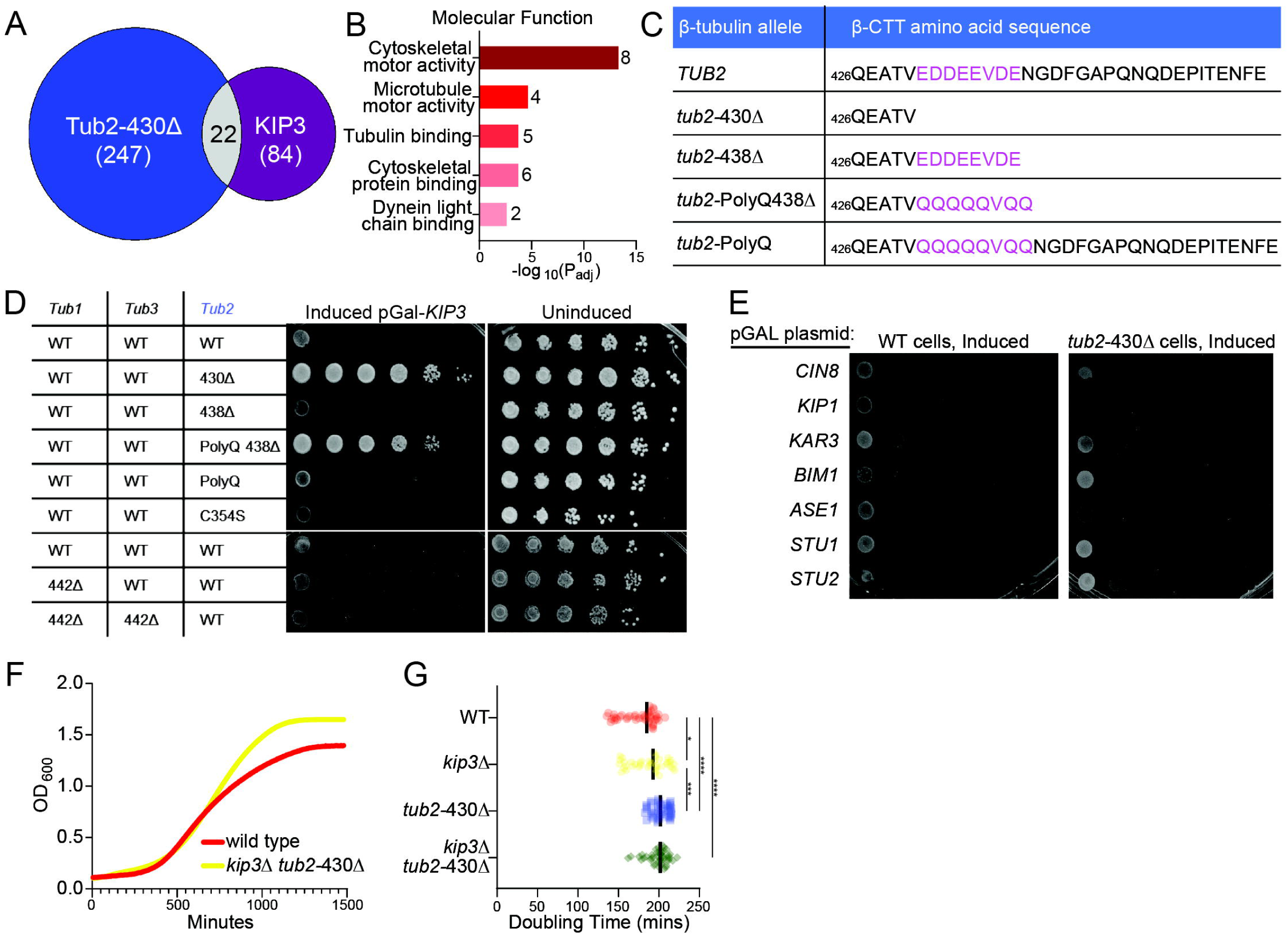
Kip3 requires β-CTT for function. **(A-B)** Results comparing negative genetic interactions between a β-CTTΔ screen and a Kip3Δ screen. **(A)** Venn diagram displaying the overlapping genes between the two screens. **(B)** The 22 overlapping genes were further sorted by gene ontology using the Gprofiler program and the top 5 molecular functions are listed. Numbers listed next to bars depict the number of genes associated with the function. **(C)** Table of genome-edited mutants that alter the composition of the β-CTT. In magenta are the 8 residues that reside in the acidic patch which contains the greatest enrichment of negatively charged residues. **(D-E)** We utilized a galactose inducible plasmid with wildtype Kip3 **(D)** or various MAPs **(E)** tagged with a triple affinity tag comprised of His6-HA^epitope^-3C^protease^ ^site^-ZZ^protein-A^ at the C-terminus. All overexpression strains were grown to saturation and a tenfold dilution series were spotted onto inducing or noninducing medium and grown at 30°C. **(D)** Kip3 overexpression into the β-CTT panel. **(E)** Overexpression of MAPs in wild-type or *tub*2-430Δ cells. **(F)** Example growth curve from a liquid growth assay of wildtype cells completed in a Cytation3 plate reader over the span of 24 hours where OD600 measurements were taken every five minutes. **(G)** Doubling time calculated from growth curves for wild-type, *kip*3Δ, *tub*2-430Δ, and a *kip*3Δ/*tub*2-430Δ double mutant. A one-way Anova with a Tukey’s Post-Hoc test was completed (*p=0.017, ***p=0.0003, ****p<0.0001, n ≥ 42). Bars represent the median value for each genotype.

We conducted two experiments to test whether β-CTT is required for Kip3 function. First, we conditionally overexpressed Kip3 in yeast cells with various β-tubulin tail mutations to identify regions of the tail that are required for Kip3 activity (Figure 1C). In wild-type cells, overexpression of Kip3 inhibits cell proliferation presumably by creating high levels of MT depolymerase activity (Figure 1D). We predicted that ablating regions of β-CTT that are required for Kip3 function should rescue proliferation during Kip3 overexpression. Consistent with our prediction, *tub*2-430Δ mutant cells lacking β-CTT rescue growth inhibition by Kip3 overexpression (Figure 1D). Additionally, *tub*2 mutants where the negative charges of β-CTT are neutralized (PolyQ), rescue Kip3 overexpression (Figure 1C and D). These results demonstrate that the negatively charged amino acids within β-CTT are necessary for Kip3 activity. This effect does not appear to be an indirect consequence of stabilizing MTs, since cells with the *tub2*-C354S mutation, which stabilizes MTs^22^, are sensitive to Kip3 overexpression (Figure 1D). Furthermore, this effect is specific to β-CTT; cells lacking α-CTT are sensitive to Kip3 overexpression, similar to wild-type controls (Figure 1D). To test whether the requirement for β-CTT is specific for Kip3, we conducted separate experiments to overexpress other MAPs involved in controlling MT dynamics in both wild-type and *tub*2-430Δ cells. Overexpression of these MAPs, including kinesin-5 and kinesin-14 motors, inhibits the proliferation of both wild-type cells and *tub*2-430Δ mutants (Figure 1E). This suggests that the requirement for β-CTT is specific to Kip3.

For our second experiment, we designed epistasis experiments to test the prediction that if Kip3 is non-functional in *tub*2-430Δ mutant cells, then combining a Kip3 null mutant (*kip*3Δ) with the *tub*2-430Δ mutant should not create additive phenotypes. We previously found that *tub*2-430Δ mutants exhibit a longer doubling time due to defects in mitotic spindle assembly and delay by the spindle assembly checkpoint^23^. As predicted, we find that *tub*2-430Δ and *kip*3Δ single mutant strains exhibit a significantly longer doubling time than wild-type cells (p<0.0001, p=0.017), and the *tub*2-430Δ *kip*3Δ double mutant strains exhibit a doubling time that is indistinguishable from the *tub*2-430Δ single mutant. (p=0.8087, Figure 1F and G). We also examined anaphase spindle morphology in single versus double mutants. Consistent with previous reports, *kip*3Δ single mutants exhibit over-elongated anaphase spindles, which may be attributed to failure to regulate interpolar MT length in late mitosis (Figure S1C and D)^24^. *tub*2-430Δ *kip*3Δ double mutant cells also exhibit over-elongated anaphase spindles (Figure S1C and D). These epistasis results suggest that Kip3 requires β-CTT for most, but perhaps not all functions.

### β-CTT is necessary to promote Kip3’s depolymerase activity

Previous studies have reported multiple activities for Kip3, including plus-end directed motility, depolymerase activity, and MT crosslinking^5,13,25^. It is unclear whether β-CTT is necessary for one or more of these activities. To test depolymerase activity, we transformed the Kip3 overexpression plasmid into wild-type or *tub*2-430Δ cells expressing GFP fused to an extra copy of α-tubulin (GFP-Tub1) to visualize MTs (Figure S2). In wild-type cells, the number of MTs decreased after one-hour of Kip3 overexpression (Figure 2A and B). After three hours of Kip3 overexpression, we observed the appearance of blob-like accumulations of GFP-Tub1 in the cytoplasm (Figure 2A, S3A). We never observed these accumulations in control cells with normal levels of Kip3 (data not shown). In separate experiments, we find that Kip3 co-localizes to tubulin accumulations (Figure S3B). This suggests that Kip3 sequesters tubulin when off the MT. We do not see the same tubulin accumulations when we overexpress other MAPs in wild-type cells expressing GFP-Tub1 (Figure S3C), indicating that tubulin accumulations are specifically created by high levels of Kip3 (Figure S2). We find that wild-type cells lose approximately 50% of MT structures over time of Kip3 overexpression (Figure 2B). When Kip3 is overexpressed in *tub*2-430Δ cells, MTs persist at a higher frequency than in wild-type cells even after three hours of Kip3 overexpression and there is a lower frequency of the tubulin accumulations over time (Figure 2A and B, S3A). Importantly, total tubulin levels are similar in wild-type and *tub2*-430Δ cells, and do not change during Kip3 overexpression (Figure S2C and D). These results suggest that Kip3 requires β-CTT for its depolymerase activity.

**Figure 2:**
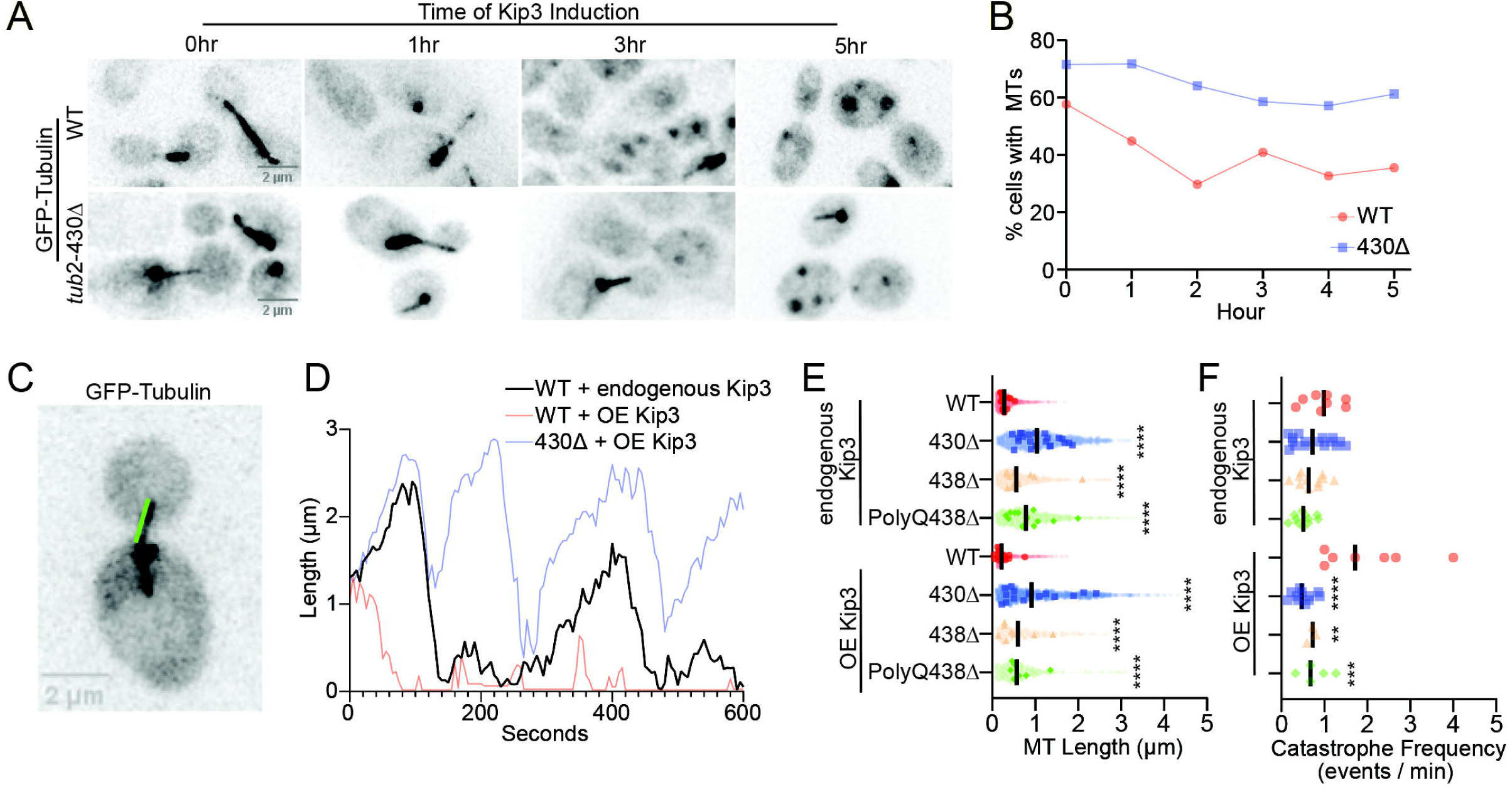
β-CTT is necessary to promote Kip3’s depolymerase activity. **(A)** Representative images of wild-type and *tub*2-430Δ cells expressing GFP-Tub1 fixed between 0-5hours after Kip3 induction. **(B)** Percent of cells with MT structures of fixed cells quantified between 0-5hrs. **(C-F)** Cells were induced for Kip3 overexpression for one-hour and imaged via confocal microscopy every 5 seconds for 10 minutes. For all MT dynamic analyses each genotype was analyzed in addition to an empty plasmid control **(C)** Representative image of a wild-type cell expressing GFP-tubulin. Green line demonstrates the measured MT length between the plus end and the spindle pole body. **(D)** Example life plot of MT length over time from of a single MT for a wild-type control MT, and MTs with Kip3 overexpression from a wild-type and a *tub*2-430Δ cell. **(E)** Quantification of all MT lengths for the panel of β-tubulin mutants overlapped with median MT lengths for individual MTs (n ≥ 440). **(F)** Quantification of catastrophe frequency. Each dot represents measurements from MT lengths from individual cells. **(E-F)** A one-way Anova with a Tukey’s Post-Hoc test was completed for all Endogenous Kip3 genotypes or Kip3 overexpression separately (****p<0.0001, ***p=0.0004, ** p=0.0023).

We next compared MT dynamics in wild-type and β-CTT mutants, either with normal or overexpressed Kip3. To test this, we conducted live cell imaging of cells expressing GFP-Tub1, measuring the lengths of individual astral MTs over a ten-minute period (Figure 2C and D)^26^. With normal levels of Kip3, wild-type cells exhibit a lower median MT length than all *tub*2 mutants we tested (0.28 µm). *tub*2-430Δ cells displayed the longest median length (1.04 µm), followed by *tub*2-PolyQ-438Δ (0.78 µm), which aligns with previously published results (Figure 2E)^27^. *tub*2-438Δ showed an intermediate phenotype, significantly longer than wild type but significantly shorter than *tub2-430Δ* (0.56 µm, Figure 1C, 2E). We also measured the frequency of catastrophes, which are created by Kip3 depolymerase activity. With normal Kip3 expression levels, wild-type cells exhibit more catastrophes per minute than all β-CTT mutants tested, but none of these differences were significant (Figure 2F).

We predicted that slightly overexpressing Kip3 in wild-type cells would increase catastrophe frequency due to increased depolymerase activity at plus ends, and lead to shorter median MT lengths. Indeed, overexpressing Kip3 for one hour in wild-type cells decreases median MT length (0.22 µm) and increases catastrophe frequency (WT control vs OE, p=0.0007; Figure 2E and 2F). In contrast, overexpressing Kip3 in *tub*2-430Δ cells does not change median MT lengths or catastrophe frequency significantly (0.92µm; Figure 2E and F). This suggests that β-CTT is necessary for Kip3 depolymerase activity. For the other β-CTT mutant cells, Kip3 overexpressed *tub*2-438Δ cells had no change in median MT length (0.60 µm) while *tub*2-PolyQ-438Δ had a slight decrease in median MT length (0.57 µm; Figure 2F). Both *tub*2-438Δ and *tub*2-PolyQ-438Δ had trending increases in catastrophe frequency though not significant (Figure 2F). The lack of changes in the other β-CTT mutants suggests that the charge of the entire tail, not just the acidic patch, may be necessary for Kip3 depolymerase activity.

### β-CTT decreases Kip3 localization at the plus end of the microtubule

Loss of Kip3 depolymerase activity in cells lacking β-CTT could be attributed to disruption of Kip3 binding to MTs, plus-end directed motility, or the depolymerase mechanism. To measure MT binding and plus-end accumulation, we imaged Kip3 on MTs by collecting timelapse, z-series images in wild-type, *tub*2-430Δ, *tub*2-438Δ, and *tub2*-C354S cells expressing Kip3-mNeonGreen and mRuby-Tub1. We focused on cells in preanaphase, when single astral MTs extend from the short mitotic spindle (Figure 3A). We identified polymerizing MTs from the timelapse and quantified the cumulative intensity of Kip3-mNeonGreen along the entire MT by taking line scans starting from the MT plus end to the cytoplasmic face of the SPB. We then divided total Kip3-mNeonGreen intensity by the MT length to measure Kip3 intensity per µm (Figure 3B, S4A and B). *tub*2-430Δ cells show, on average, increased Kip3-mNeonGreen accumulation on MTs compared to wild-type cells, while *tub*2-438Δ cells show no significant difference compared to wild-type (Figure 3B). The *tub2*-C354S mutant, which stabilizes MTs^22^, is also similar to wild-type controls in this experiment, indicating that the increased Kip3 on *tub2*-430Δ MTs is not an indirect consequence of longer MTs (Figure 3B, S4A). These results show that β-CTT is not required for Kip3 to bind MTs.

**Figure 3:**
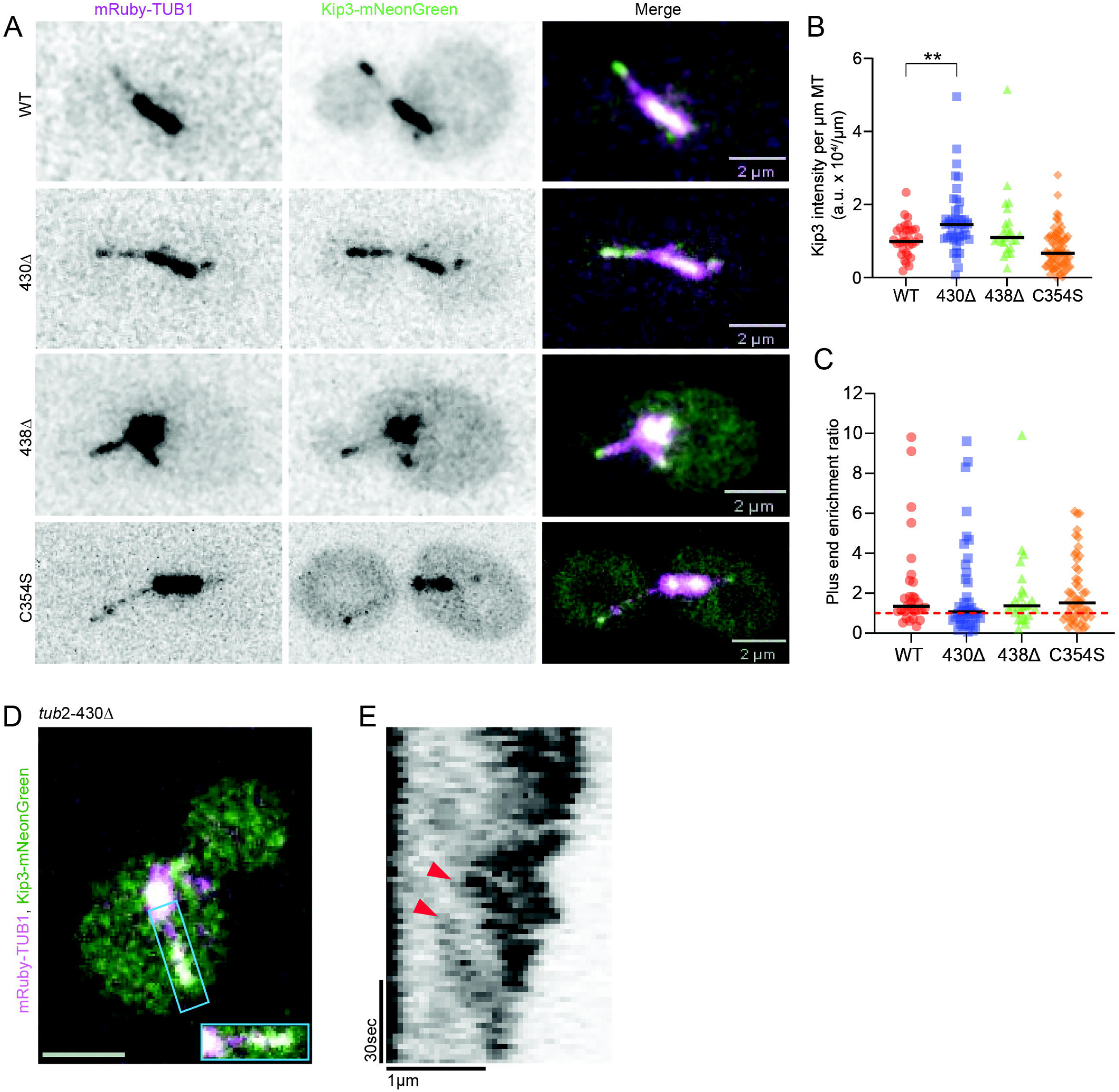
β-CTT decreases Kip3 localization at the plus end of the microtubule. **(A)** Representative images of wild-type, *tub*2-430Δ, *tub*2-438Δ, and *tub*2-C354S strains tagged with mRuby-TUB1 to visualize MTs and Kip3-mNeonGreen. **(B)** Line scan analysis of Kip3-mNeonGreen intensity across the entire MT lattice divided by the length of the MT. **(C)** Ratio of Kip3-mNeonGreen intensity at the plus end versus the middle of the MT. Red dotted line = 1 **(B-C)** A one-way Anova with a Tukey’s Post-Hoc test was completed where p<0.05 is significant. Bars represent the median value. (**p=0.0046, n ≥ 30). Each dot represents a MT from an individual cell. **(D-E)** A PDR mutation was added in the *tub*2-430Δ mutant and cells MTs were stabilized with the drug epothilone A. **(D)** Representative image of a treated cell **(E)** Kymograph of Kip3-mNeonGreen along the MT lattice taken from a 2-minute time lapse video with 2 second intervals. Arrows depict the beginning of a motility event.

The increased accumulation of Kip3 on MTs in *tub*2-430Δ cells raises the question of whether β-CTT is necessary for Kip3 plus-end directed motility. Motility should create an enrichment of Kip3 at the plus end, compared to the lattice of the MT. To test this prediction, we compared the intensity of Kip3-mNeonGreen in a 413 nm^2^ (7×7 pixels; see Methods) region at the plus end of the MT to an equally sized region at the geometric center of the MT. We used these measurements to create plus-end enrichment ratios by taking the sum intensity of Kip3-mNeonGreen signal at the MT plus end over the sum intensity at the MT center for each MT (Figure S4C and D). Ratios > 1 indicate plus-end accumulation, while ratios ≤ 1 indicate lack of plus-end accumulation. We validated this method using astral MTs labeled with GFP-Tub1, which are not expected to exhibit plus-end enrichment; indeed, these show a median ratio of 0.92 (plus end/center; data not shown). *tub*2-430Δ cells exhibit significantly lower plus-end enrichment ratios compared to wild-type controls (Figure 3C; medians, WT: 1.347, 430Δ: 1.065; Fisher’s Exact Test: p<0.0001;). In contrast, *tub*2-438Δ cells exhibit Kip3 plus-end enrichment ratios that are not different from wild-type (Fisher’s Exact Test: p=0.2393, median: 1.363), while *tub*2-C354S cells exhibit a slight increase in the plus-end enrichment ratio (Figure 3C; Fisher’s Exact Test: p=0.0114, median, 1.514). These results indicate that β-CTT may be important for plus-end directed motility.

### β-CTT inhibits Kip3 lattice binding and alters motility

To assess Kip3 motility, we first measured Kip3 dynamics in living cells. Since astral MTs in budding yeast are typically short (Figure 2E), we created longer MTs by treating cells with epothilone A which stabilizes MTs. We collected timelapse images at 2-second intervals in cells expressing Kip3-mNeonGreen and mRuby-Tub1 and used image series to generate kymographs of Kip3-mNeonGreen movement on stable MTs (Figure 3D and E). We observed multiple motility events of Kip3 moving towards the plus end of the MT in *tub*2-430Δ (arrowheads, Figure 3E). This suggests that β-CTT is not absolutely required for Kip3 motility.

To carefully compare the rate of Kip3 motility on wild-type MTs vs MTs lacking β-CTT, we reconstituted motility *in vitro* from yeast whole cell lysates and total internal reflection fluorescence (TIRF) microscopy. Previous studies have shown that lysates from yeast cells arrested at metaphase grow dynamic microtubules and that Kip3 bound to these MTs displays movement towards the plus end^28^. To reconstitute Kip3 motility, we assembled hybrid MTs where GMPCPP-stabilized, rhodamine-labeled porcine brain tubulin MTs seeded the assembly of yeast tubulin from a high concentration of metaphase-arrested yeast cell lysate (20 mg/ml) that were stabilized by adding epothilone A (2 µM; see Materials and Methods). After incubating at 30°C for 10 minutes, imaging chambers were washed with a high salt buffer (750 mM KCl) to remove MT-bound MAPs while maintaining stable MTs with epothilone A, followed by flowing in a lower concentration of the same whole cell lysate (9 mg/ml) and still maintaining stable MTs with epothilone A (100 µM; Figure 4A). We measured the lengths of unlabeled yeast MTs using Interference Reflection Microscopy (IRM) after flowing in our lower concentration and find that MTs are stable and that wild-type and *tub*2-430Δ MT lengths are on average 11.10 ± 2.44µm and 13.7 ± 1.25µm, respectively (mean ± SD; data not shown). Similar to our results in cells, we find that the amount of Kip3-mNeonGreen bound to *tub*2-430Δ MTs is increased compared to wild-type MTs (Figure 3, 4B). To quantify this difference, we used line scan analyses to measure the Kip3-mNeonGreen intensity on the GMPCPP seed and on the adjacent yeast MT lattice. As expected, the amount of Kip3 bound to GMPCPP seeds did not change between the wild-type and *tub2*-430Δ extracts (Figure 4C; p=0.3844). However, the amount Kip3 bound to *tub*2-430Δ MT lattice is significantly greater than wild-type MT lattice (Figure 4D; p=0.0337). These results confirm that β-CTT weakens the affinity of Kip3 for the MT lattice.

**Figure 4:**
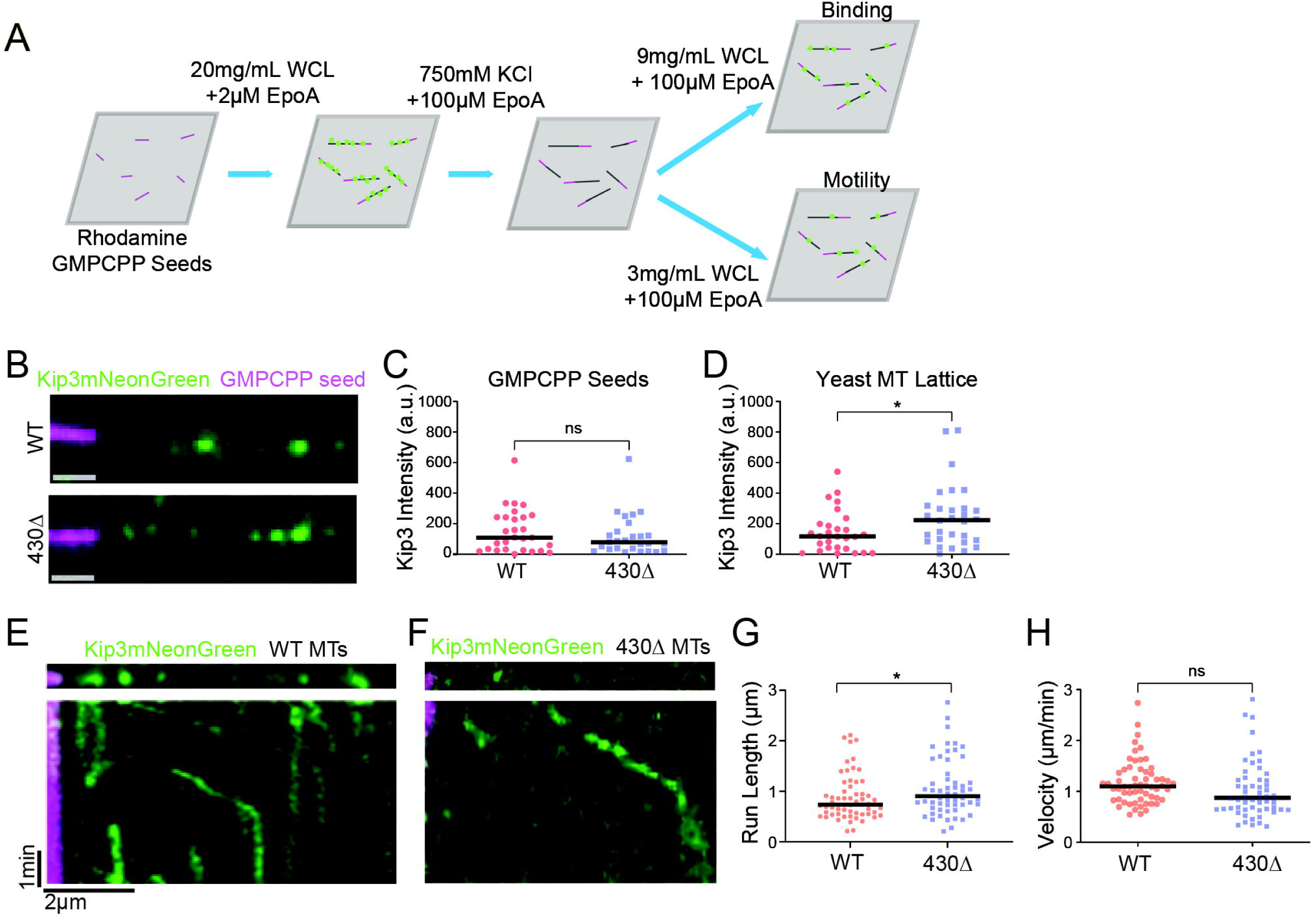
β-CTT inhibits Kip3 lattice binding and alters motility. **(A)** Diagram of *in vitro* WCL-TIRF reaction set-up. A high concentration of lysate was added into a chamber with rhodamine labeled GMPCPP seeds to grow MTs. MTs were stabilized with epothilone A (EpoA) and high salt washes were conducted to remove proteins from MTs. Finally, a low concentration of lysate was added to visualize individual Kip3-mNeonGreen for binding and motility events. **(B)** Example images of Kip3-mNeonGreen binding along unlabeled wild-type or *tub*2-430Δ MTs. **(C-D)** Quantification of Kip3-mNeonGreen binding 1µm into the GMPCPP seeds or 1µm into the MT grown off of the same seed (*p=0.0337, n ≥ 28). **(E-F)** Example kymographs videos imaged every 5 seconds for 5 minutes for wild-type and *tub*2-430Δ MTs. **(G)** Quantification of Kip3 run length (*p=0.0454, (n ≥ 56). **(H)** Quantification of Kip3 velocity from each run (n ≥ 56). A two-tailed T-test was conducted for each quantification where p<0.05 is significant. Bars represent the median value. Each dot represents individual Kip3 foci.

We observed plus-end directed movement of individual Kip3 foci on wild-type MTs in our experiment with 9 mg/mL of cell lysate (Figure 4E). In contrast, *tub2-*430Δ MTs we more crowded with Kip3 foci in 9 mg/mL lysate, which confounded the identification of motility events. We improved this by decreasing the concentration of *tub*2-430Δ lysate to 3 mg/mL, which enabled clear identification of motile Kip3 foci (Figure 4F). We generated kymographs of Kip3 motility on wild-type and *tub*2-430Δ MTs from timelapse TIRF images collected at 5 second intervals for 5 minutes (Figure 4E, 4F). These kymographs show that Kip3 run length is slightly but significantly increased on *tub*2-430Δ MTs (Figure 4G, p=0.0454), while velocity is slightly slower, but the difference is not significant (Figure 4H; p=0.2028). This confirms that β-CTT is not required for Kip3 motility but may promote shorter and slightly faster runs. Together, our results indicate that β-CTT weakens Kip3 affinity for the MT lattice and decreases processivity.

### β-CTT promotes Kip3 binding to soluble tubulin heterodimers

Given that β-CTT is required for Kip3 depolymerase activity but weakens its affinity for the lattice, we next asked whether β-CTT specifically promotes Kip3 binding to curved tubulin, a state which is present at the plus end or in solution, but not in the lattice. To measure Kip3 binding to soluble tubulin, we overexpressed epitope-tagged Kip3 in wild-type or *tub2*-430Δ cells and measured the amount of tubulin bound to Kip3 precipitated from cell lysate under MT depolymerizing conditions (see Materials and Methods). If Kip3 requires β-CTT for binding to tubulin in solution we predicted that less tubulin would bind to Kip3 precipitated from *tub2*-430Δ lysate. As predicted, we find that the amount of bound tubulin is significantly decreased in *tub2*-430Δ lysates compared to wild-type control lysate (Figure 5A and 5B, Figure S5A and B). In fact, *tub2*-430Δ tubulin bound to Kip3 was no different than tubulin bound to the beads in control strains without Kip3 overexpression (Figure 5A, S5B). This suggests that β-CTT strengthens the binding of Kip3 to soluble tubulin.

**Figure 5:**
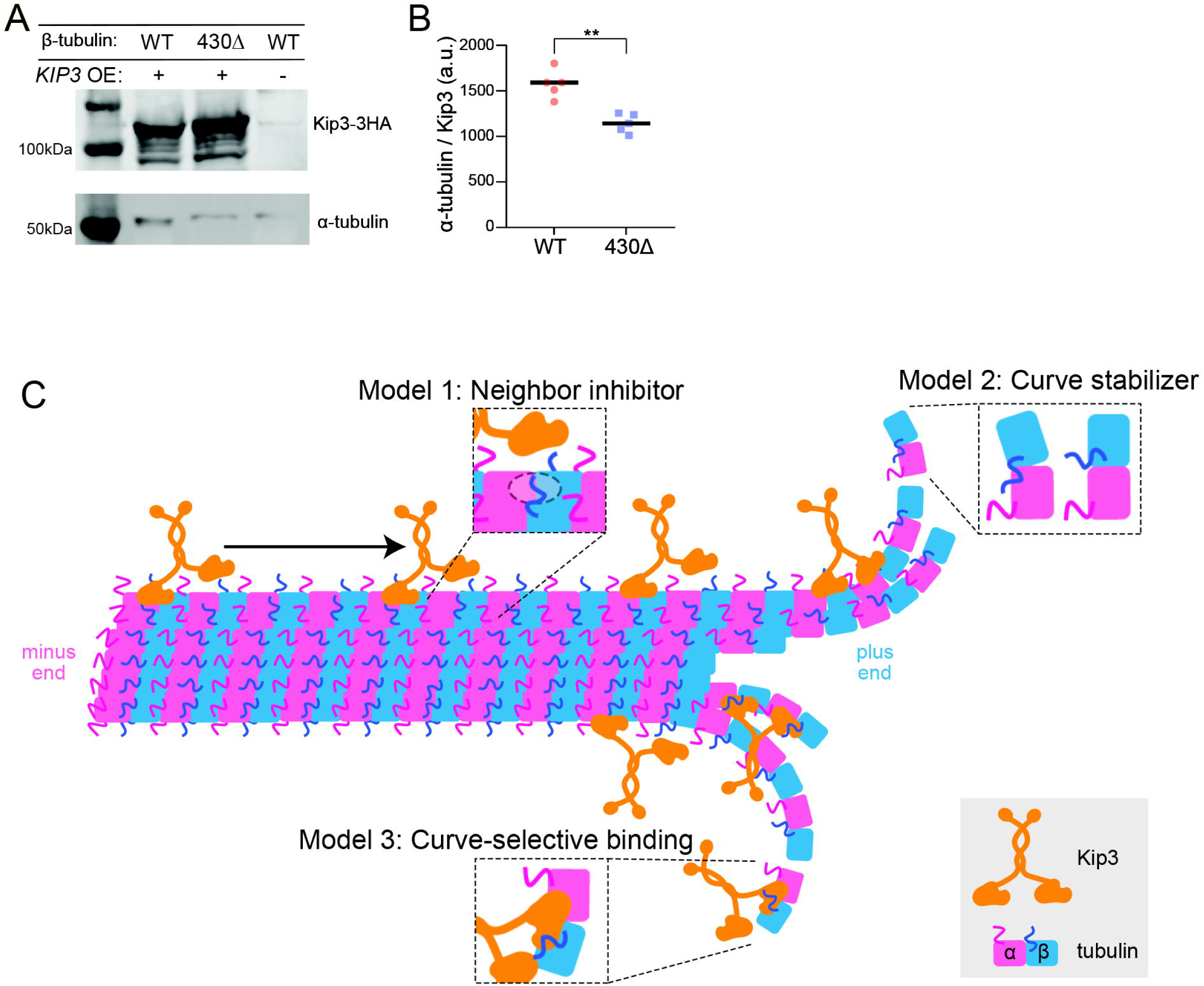
β-CTT promotes Kip3 binding to soluble tubulin heterodimers. **(A)** Example western blot probing for Kip3-3HA and α-tubulin after Kip3 pull down. **(B)** Ratio of Kip3 to tubulin bound (**p=0.0012, n = 5). **(C)** β-CTT alters Kip3 binding and velocity across the MT lattice and is necessary for Kip3 binding to free tubulin heterodimers. We propose three models of how Kip3 distinguishes between different states of tubulin through β-CTT for different activities. Model 1: β-CTT blocks Kip3 landing sites in the lattice but not at the MT plus end. Model 2: β-CTT promotes tubulin curvature at the plus end necessary for Kip3 binding and depolymerase activity. Model 3: β-CTT differentially binds to Kip3 in the lattice versus at the plus end due to changes in the Kip3 binding interface.

## Discussion

In this study, we report a novel role for the carboxy-terminal tail of β-tubulin in promoting kinesin-8/Kip3 activity. Previous studies established that Kip3 binds to MTs and uses plus-end directed motility to accumulate at plus ends where it promotes MT depolymerization^5,13^. However, it is unclear what features of tubulin or Kip3 allow for a switch from binding and motility to depolymerization. We find that β-CTT dampens Kip3 binding along the lattice (Figure 3, 4) and promotes binding to tubulin in solution (Figure 5B). Without the tail, Kip3 displays increased lattice binding and accumulation at the plus end without depolymerase activity, and a decrease in binding to soluble tubulin (Figure 2-5). Our findings therefore show that β-CTT controls Kip3 activity by allowing it to discriminate between tubulin in the MT lattice versus tubulin at the plus end or in solution.

We propose three models for how β-CTT could inhibit Kip3 binding to the lattice while selectively promoting binding to tubulin at plus ends or in solution. In the first model, β-CTT inhibits Kip3 binding to the lattice by acting in trans, with CTTs on one heterodimer docking on the neighboring heterodimer to block Kip3’s binding site (Model 1, Figure 5C). At the plus end, where protofilaments are splayed^29^, CTTs may be unable to reach across to neighboring heterodimers and unable to inhibit Kip3 binding, thereby creating a zone of strong binding and depolymerase activity. However, this model is not supported by our pull-down results, where we found that Kip3 binds more wild-type tubulin in solution, compared to *tub2*-430Δ tubulin in solution (Figure 5B). In the second model, β-CTT promotes tubulin conformational change from straight in the lattice to curved at the plus end (Model 2, Figure 5C). This model is consistent with previous work from our lab showing that MTs treated with subtilisin to proteolytically remove β-CTTs have plus ends that appear less tapered and more blunt, suggesting that β-CTT normally promotes protofilament curvature even in the absence of MAPs^27^. This work suggests that β-CTTs are important for creating the curved plus end morphology that Kip3 tightly binds to for depolymerization. While this model could explain a role for β-CTT in creating the binding state for Kip3 depolymerase activity at the plus end, it does not obviously explain how β-CTT inhibits Kip3 binding to the MT lattice. In the third model, β-CTT are repositioned when tubulin transitions from the straight conformation in the lattice to the curved conformation at the plus end and in doing so switch from a position that disrupts Kip3 binding to a new position that contributes to the binding interface and stabilizes the Kip3-curved tubulin complex that promotes depolymerization (Model 3, Figure 5C). This model is supported by previous structural evidence that the motor domain of Kip3 exhibits conformational changes on lattice tubulin vs curved tubulin oligomers^17^. Thus, it is possible that new binding sites are available in the motor when Kip3 is at the plus end for tight binding to β-CTT that are not accessible when the motor is on the lattice. More work is necessary to uncover what regions of Kip3 are interacting with β-CTT for these changes in binding.

It is important to note that our results do not eliminate the possibility of indirect effects of β-CTT loss and/or Kip3 mislocalization on other MAPs. Our previous work shows that β-CTT affects other MAPs, in addition to Kip3. Other MAPs discovered as being candidates for interacting with β-CTT from the genetic screen include the kinesin-5 motor Cin8, the MT crosslinker Ase1, and plus end binding proteins such as Bim1^10,11^. We find that the overexpression of these MAPs in cells expressing GFP-Tub1 results in abnormal spindles and MT morphology that are different from Kip3 overexpression phenotypes (Figure S3C, 2A). Overexpression of these MAPs is lethal in wild-type cells but removing β-CTT did not rescue overexpression lethality like with Kip3 (Figure 1D and E). These differences highlight the importance of differential regulation of MAPs by the tubulin CTTs. Additionally, we speculate that accumulation of Kip3 along MTs and at the plus end in β-CTT mutants (Figure 3, 4) may crowd the plus end and thus may alter the binding or activity of other regulators. More work needs to be completed to fully understand how the β-CTT identifies and differentially regulates these MAPs and motors.

As the negative interactions from the *tub*2-430Δ and *kip*3Δ genetic screens did not perfectly overlap, it is reasonable to conclude that not all Kip3 functions, as mentioned above, require interaction with β-CTT. Kip3 functions include regulating MT attachments at the cell cortex for spindle alignment at the bud neck, kinetochore clustering during metaphase, and crosslinking and sliding antiparallel MTs at the spindle midzone ^5,25,30^. Our results provide potential to separate these functions based on the dependence of Kip3 on β-CTT and robust depolymerase activity. For example, the tail domain of Kip3 is required for MT crosslinking and sliding of the spindle^25^. As we find that β-CTT is necessary for the depolymerase activity of Kip3 (Figure 2), which is mediated by the motor domain and not the tail^16^, we find no evidence that crosslinking activity requires the Kip3-β-CTT complex. Our spindle length analyses show that our *tub*2-430Δ and *kip3*Δ double mutant result in an additive effect on anaphase elongation resulting in longer spindles compared to wild-type cells or any individual mutations (Figure S1D). A more severe phenotype in the double mutant supports the idea that not all Kip3 functions are regulated by β-CTT. More work is necessary to distinguish all Kip3 activities regulated by β-CTT.

Our model for Kip3 regulation by β-CTT may also have implications for other kinesins by determining whether those motors act as transporters or plus end regulators based on relative affinities for the MT lattice vs tubulin heterodimer. For example, the kinesin-1 family transports cargo via processive motility towards the plus end of the MT stimulated by strong lattice affinity when bound by ATP^31^. At the plus end, kinesin-1 does not exhibit tight binding and instead walks off the MT, suggesting that kinesin-1 does not have high affinity for the curved microtubule ends^16,31^. On the other end of the spectrum, the kinesin-13 family members exhibit weak lattice binding and strong binding to MT ends, causing the motor to diffuse in solution to the MT ends rather than exhibit directional motility^3,32^. At the MT ends, kinesin-13 binds tightly to the terminal tubulin heterodimers and utilizes ATP hydrolysis to dissociate the dimers and cause depolymerization while remaining strongly bound to the lattice^32,33^. Kip3, and the kinesin-8 family, appear to be multi-tasking motors that can exhibit both of these behaviors. Kip3 is a highly processive motor with affinity for the MT lattice like kinesin-1 but exhibits a stronger affinity for tubulin at the plus end^16^, like kinesin-13. Therefore, Kip3 can walk along the lattice, but also dwell at the plus ends, where it binds tightly to terminal tubulin subunits to destabilize tubulin binding to other tubulins in the MT. Unlike kinesin-13, the ATPase activity of Kip3 at the plus end is not necessary for Kip3 induced depolymerization^16^. This may be attributed to Kip3 falling off of the lattice with the bound heterodimer^13^ rather than the ability to stay bound and continue to depolymerize like kinesin-13. We postulate that β-CTT is key to Kip3’s multi-tasking behavior by enabling the switch into a free-tubulin binding mode at the plus end.

## Materials and Methods

### Yeast strains and manipulations

All experiments were done in *Saccharomyces cerevisiae* with the S288C genetic background. The *kip*3Δ deletion mutant was generated by conventional PCR-based methods and insertion of a URA marker at the genomic locus^34^. All β-tubulin tail mutants were made at the native, chromosomal *TUB*2 locus by PCR-mediated mutagenesis^23,27^. The *KIP*3 overexpression plasmid and empty plasmid control used for microtubule dynamics and co-immunoprecipitation experiments is from the Yeast ORF collection (pJM214, pJM294; Gelperin et. al, 2005). GFP-Tub1 (pSK1050)^35^ or mRuby-TUB1 (pcDNA3:*mRuby*2)^36^ were integrated into the yeast genome and integrated adjacent to the *TUB*1 locus to visualize microtubules. The mNeonGreen fluorescent protein^37^ was fused to the carboxy-terminus of *KIP*3 at the native locus via PCR amplification from pJM261 using oligos #80 and 81. *Cdc*-23 alleles were integrated by conventional PCR-based methods and then crossed to strains containing the Kip3-mNeonGreen tag for reconstitution experiments. A detailed list of all strains, plasmids, oligos, antibodies, chemicals, and softwares used in this study can be found in the Key Resource Table below.

### Gene ontology

Genetic screen data from cells lacking the β-CTT (*tub*2-430Δ) or Kip3 (*kip3*Δ) was data from Aiken et al.and Costanzo et al., respectively^10,21^. Negative interactions from each list were included with stringent cutoff: score <-0.12, p-value < 0.05. Any genes included in the *kip*Δ screen that were excluded from the β-CTT screen were removed. All genes for the β-CTT were imputed into GeneMANIA. *KIP*3 was identified as highly related to this network for biological, molecular, and cellular processes. Venn diagram of genes found as negative interactions between *tub*2-430Δ and *kip3*Δ screens was generated using the Venn Diagram Generator (http://jura.wi.mit.edu/bioc/tools/venn.php). GO-terms for overlapping negative interactors between each screen was analyzed using G-profiler (Tartu, Estonia)^38^.

### Kip3 overexpression

A galactose inducible plasmid with wildtype Kip3 tagged with a triple affinity tag comprised of His6 -HA^epitope^ - 3C^protease^ ^site^ -ZZ^protein-A^ at the C-terminal end (pJM214) or an empty plasmid control (pJM294) was transformed into wildtype yeast strains or strains with β-tubulin tail mutants. Cells were grown in non-inducing media (2% raffinose and CSM-URA; Sunrise Science; Knoxville, TN) until log phase before galactose was added to 2% at 30°C with shaking for up to 5 hours. See ‘microtubule dynamics analysis’ and ‘yeast fixation’ for further methods.

### Liquid growth assay

Yeast cells were grown to saturation in 3 ml of rich media (2% glucose, 2% peptone, and 1% yeast extract) at 30°C and diluted 500-fold into fresh media. 200µl of each diluted cultures were transferred to a 96-well plate and incubated at 30°C in a Cytation3 plate reader (BioTek; Winnoski, VT) with single orbital shaking. The OD_600_ was measured every 5min for 24hrs. Doubling time was calculated for each strain by fitting growth curves to a nonlinear exponential growth curve using MATLAB code from Fees et al, 2018^9^.

### Solid growth assays

Tubulin-CTT mutants with the wildtype Kip3 overexpression (pJM 214) and other MAP overexpression (pJM207-212, 659) were grown to saturation at 30°C in non-inducing media (2% raffinose and CSM-URA) and a 10-fold dilution series of each was spotted to either non-inducing (2%glucose, 2% raffinose, and CSM-URA) or inducing (2,% galactose, 2% raffinose, and CSM-URA) plates. Plates were grown at 30°C for 3 days and then imaged.

### Live-cell imaging

Yeast cells were grown overnight at 30°C in rich media, then diluted in synthetic media (2% glucose and CSM; Sunrise Science; Knoxville, TN) and grown until log phase at 30°C. Cells were adhered in flow chambers with concanavalin A and sealed with VALAP (equal parts Vaseline, lanolin, and paraffin)^36^. For all imaging experiments, the stage of the scope remained 30°C. Images were collected on a Nikon Ti-E spinning disk confocal microscope with a 100x, 1.45 NA CFI Plan Apo oil emersion objective, confocal scanner (CSU10; Yokogawa), piezoelectric stage (Physik Instrumente; Auburn, MA), 488-nm and 561-nm lasers (Agilent Technologies; Santa Clara, CA), and an EMCCD camera (iXon Ultra 897; Andor Technology, Belfast UK) using NIS Elements software (Nikon; Tokyo, Japan). Stage was heated to 30°C using an ASI 400 Air Stream Incubator (NEVTEK; Williamsville, VA).

### Microtubule and spindle dynamics acquisition and analysis

Yeast cells were grown as described in ‘live-cell imaging’ or for Kip3 overexpression, cells were grown as described in ‘Kip3 overexpression’ and induced with 2% galactose to image over the 1-hour time point. For all microtubule dynamics acquisitions, time-lapse images were taken every 5s for 10min with a z-series of 6µm separated by 500nm steps. Z-series images were max-projected and the lengths of astral microtubules for each timepoint was measured using the line scan tool in FIJI/ImageJ (Wayne Rasband, NIH)^39^. Microtubule length changes over time was used to calculate catastrophe frequency using a custom MATLAB code^26^. Polymerization and depolymerization events were defined as length increases or decreases by at least 0.5 µM across a minimum of 3 timepoints and fitting a linear regression with an R-value<0 and R^2^≥0.9. For anaphase spindle analyses, time-lapse images were taken every 10s for 5min. Z-series images were max-projected, and the length of the spindle was measured using the line scan tool in FIJI/ImageJ.

### Yeast fixation and MT analysis

Wildtype or *tub*2-430Δ cells with GFP-Tub1 and the Kip3 overexpression plasmid were grown in non-inducing media (2% raffinose and CSM-URA), then induced with 2% galactose for 0-5 hours and fixed. To fix 4-parts cell suspension was added to 1-part 5X fix solution (0.5 M K_3_PO_4_, pH 7.0; 18.5% formaldehyde) and incubated for 5min. Cells were pelleted, and supernatant removed. Cells were washed once with quencher solution (0.1 M K_3_PO_4_, pH 7.0; 0.1% Triton; 10mM ethanolamine), once with 0.1 M K_3_PO_4_, pH 7.0 to the desired volume. Cells were imaged on the confocal microscope. Cells were categorized as having a spindle, astral microtubules, interphase microtubules, or tubulin accumulations.

### Western blotting

To measure Kip3 protein levels, wild-type or *tub*2-430Δ cells containing the Kip3 overexpression plasmid were first grown at 30°C in 5 ml of selective media, diluted, and grown for 3 hrs before inducing overexpression with 2% galactose for 0-5 hrs. To make soluble protein lysates, cells were pelleted and resuspended in 2 M Lithium acetate and incubated at room temperature for 5min. Cells were pelleted and resuspended in 0.4 M NaOH and incubated on ice for 5 min. Cells were pelleted and resuspended in 2.5X Laemmli buffer and boiled for 5min. Before loading gels, samples were boiled and centrifuged at 6,000xg for 3min^40^.

15 µl of each sample were loaded in each lane on a 10% Bis-Tris PAGE gel in MOPs running buffer (50 mM MOPS, 50 mM TrisBase, 0.1% SDS, 1 mM EDTA, pH7.7) at 0.04 mAmps per gel for 1hr. Gels were transferred to PVDF membrane in transfer buffer (25 mM Bicine, 25 mM Bis-Tris, 1 mM EDTA, pH7.2) at 0.33 mAmps for 1hr then blocked at room temperature for 1 hr in PBS blocking buffer. Membranes were probed with primary mouse anti-6x-His (4A12E4; at 1:1000; Invitrogen; Carlsbad, Ca), mouse anti-α-tubulin (4A1; at 1:100: Piperno and fuller, 1985), mouse-anti-HA (F-7; at 1:500, Santa Cruz; Dallas, Tx), or rabbit-anti-Zwf1 (Glucose-6-phosphate dehydrogenase; at 1:10,000; Sigma; St. Louis, MO) overnight at 4°C. After incubation membranes were washed with PBS and probed in secondary goat-anti-mouse-680 (at 1:15000; LI-COR Biosciences; Lincoln, NE) and goat-anti-rabbit-800 (at 1,15,000; LI-COR) for 1 hour at room temperature. Membranes were washed twice with PBST and once in PBS and imaged on an Odyssey CLx imager (LI-COR). Bands were quantified using Image Studio (LI-COR) with Zwf1 as a loading control.

### Kip3-mNeonGreen fluorescence intensity acquisition and analysis

Cells expressing Kip3-mNeonGreen/mRuby-Tub1 in wildtype and β-tubulin mutants (yJM4867, 4868, 4870, 4986-4988, 5038, 5039) were grown and imaged as stated in ‘live-cell imaging’, using 488-nm and 561-nm lasers. Time-lapse images were taken every 5s for 1min of budding cells with a z-series of 6 µm separated by 250 nm steps, to determine if microtubules were polymerizing or depolymerizing. The first image of videos containing polymerizing astral microtubules, determined by mRuby-Tub1, were used for further analyses. Z-series images were sum-protected and Kip3 intensity was determined through line scan analyses from the spindle pole body to the microtubule plus end in FIJI/ImageJ (Wayne Rasband, NIH)^39^. To measure Kip3 intensity at the middle of the microtubule, line scans across the mRuby-Tub1 signal were used to determine the midpoint of the microtubule and a 7-by-7-pixel box was drawn to measure intensity. To determine the intensity of Kip3 at the microtubule plus end, a 7-by-7-pixel box was drawn around the center of the microtubule plus end, determined by the mRuby-Tub1 signal. The per-pixel cytoplasmic background from each cell was determined with a 7-by-7-pixel box, normalized to area, and subtracted from each measurement. The ratio of Kip3 at the plus end to the middle of the microtubule was calculated by dividing the normalized intensities of Kip3 at the plus end to Kip3 at the middle of the microtubule. A minimum of 30 cells were analyzed across each strain.

### Whole cell lysate *in vitro* binding and motility assays

Strains with c*dc*-23 allele (yJM5333, 5335) were grown overnight at 25°C to OD_600_∼0.40 in rich media then shifted to 37°C for 3hrs to arrest before being harvested. Cells were pelleted, washed with water, and diluted by 10xBRB80 (800mM PIPES, 10mM MgCl_2_, 10mM EGTA; pH6.9) with 150mM KCl. Yeast slurries were flash frozen in liquid nitrogen and stored at −80°C. To generate whole cell lysates, frozen cells were lysed by grinding in a Mixer Mill MM 400 (Retsch; Hamburg, Germany) for two 110s cycles at 30Hz. Powdered lysate was stored at −80°C.

Protocol to generate microtubules and track Kip3 movement was based on Bergman et al., 2019. To prep lysates, yeast powder was diluted for low and high concentrations by adding 2 µl or 0.1 µl of 10xBRB80 plus 150 mM KCl per 1 mg of powder respectively. Lysates were clarified by centrifugation at 100,000xg for 30 min at 4°C. Supernatants were removed and protein concentration was determined by a Quick Start Bradford Protein Assay (BioRad, Hercules; CA). To polymerize microtubules a 50 µl reaction was assembled consisting of 10 µl or the high concentration of clarified lysate in 1xBRB80 plus 1 mM MgCl_2_, 8 mM ADP, 1 mM GTP, and 2 µM epothilone A and flowed into the reaction chamber. Reaction was placed on the scope at 30°C for 5-10 min until polymerization was confirmed by IRM. To remove high concentrations of MAPs, the chamber was washed twice with 50 µl of 750 mM KCl with 100 µM epothilone A for 3 min each. A 50 µl reaction was prepared for imaging using the low concentration of clarified lysate in 1xBRB80 plus 1 mM MgCl_2_, 2 mM ATP, 1 mM GTP, and 100 µM epothilone A and flowed into the reaction chamber and sealed with VALAP. The low concentration of clarified lysate was using in the low reaction at 450 ng per 50 µl for binding experiments and 150 ng per 50 µl reaction for *tub2*-430Δ motility experiments.

All images were collected at 30°C on a Ti-E microscope with a 100x, 1.49NA (CFI1160) Apochromat objective, TIRF illuminator, OBIS 488-nm and Sapphire 561-nm lasers (Coherent; Santa Clara, CA), W-View GEMINI image splitting optics (Hamamatsu Photonics; Hamamatsu, Japan), and an ORCA-Flash 4.0LT scientific complementary metal-oxide-semiconductor camera (Hamamatsu Photonics) using NIS Elements Software (Nikon). The stage was heated to 30°C using an ASI 400 Air Stream Incubator (NEVTEK). Time-lapse images were acquired using single-plane TIRF at 5 s intervals. Kip3-mNeonGreen intensity was determined through line scan analyses of 1 µm along the microtubule from the edge of the seed out towards the plus end and 1 µm into the seed in FIJI/ImageJ. The per-pixel cytoplasmic background was determined by drawing a 1µm line and measuring intensity directly next to each individual microtubule, normalized to area, and subtracted from each measurement. To analyze Kip3 motility, kymographs were generated by drawing a 10-pixel wide line along a microtubule and using the kymograph function in FIJI/ImageJ. Kymographs were analyzed by marking the first and last points of a motility event and calculating the displacement and duration between the two points. A two-tailed T-test was conducted for each quantification from 3 separate days and at least 30 microtubules where p<0.05 is significant.

### Kip3 pull-down assays

Kip3 was overexpressed in wild-type yeast strains, *tub*2-430*Δ* mutant, and an empty plasmid control strain as described in ‘Kip3 overexpression’ until saturation in 1-liter volumes before induction with galactose for 2 hrs. Yeast slurries were flash frozen in liquid nitrogen and stored at −80°C. To generate whole cell lysates, frozen cells were lysed by grinding in a Mixer Mill MM 400 (Retsch; Hamburg, Germany) for two 110 s cycles at 30 Hz. Powdered lysate was stored at −80°C. Powdered lysates were measured and 100 µl of Lysis Buffer (50 mM HEPES pH7.5, 100 mM NaOAc, 5 mM MgOAc, 0.25% NP-40, 1 mM PMSF, protease inhibitor cocktail tablet) was added for every 0.1 g of cells and vortexed. Cells were centrifuged for 5 min at ∼17,000xg and the supernatant was transferred to a new tube for whole cell lysates. Lysates were normalized to 30 mg/ml in lysis buffer. For each normalized sample, 10 µl of His-Tag Dynabeads (Thermo Fisher Scientific; Waltham, MA) (prewashed with lysis buffer) were added and placed on a rotator at 4°C for 2 hrs. Samples were washed 3x in wash buffer (50 mM HEPES pH7.5, 100 mM NaOAc, 5 mM MgOAc, 0.25% NP-40, 1 mM PMSF). Supernatant was removed from the beads and proteins were eluted off the beads by boiling in 1x SDS sample buffer for 10 min. Samples were vortexed and spun at ∼17,000xg for 1 min and the supernatant was collected for SDS-page analysis. Kip3 and tubulin were probed using anti-HA and anti-α-tubulin antibodies. A two-tailed T-test, or a one-way Anova was conducted for the Kip3 and tubulin quantification where p<0.05 is significant.

### Statistical Analysis

All experiments were conducted with two biological replicates when applicable and repeated on three different days for technical replicates. Image analysis was conducted with Fiji/ImageJ software. Significance between the experimental conditions and controls will be determined using a two-tailed T-test to calculate P-values in Prism software (GraphPad; La Jolla, CA) where P<0.05 is considered statistically significant. A One-way ANOVA with Tukey’s post-hoc test was used to compare multiple conditions.

## Key Resource Table

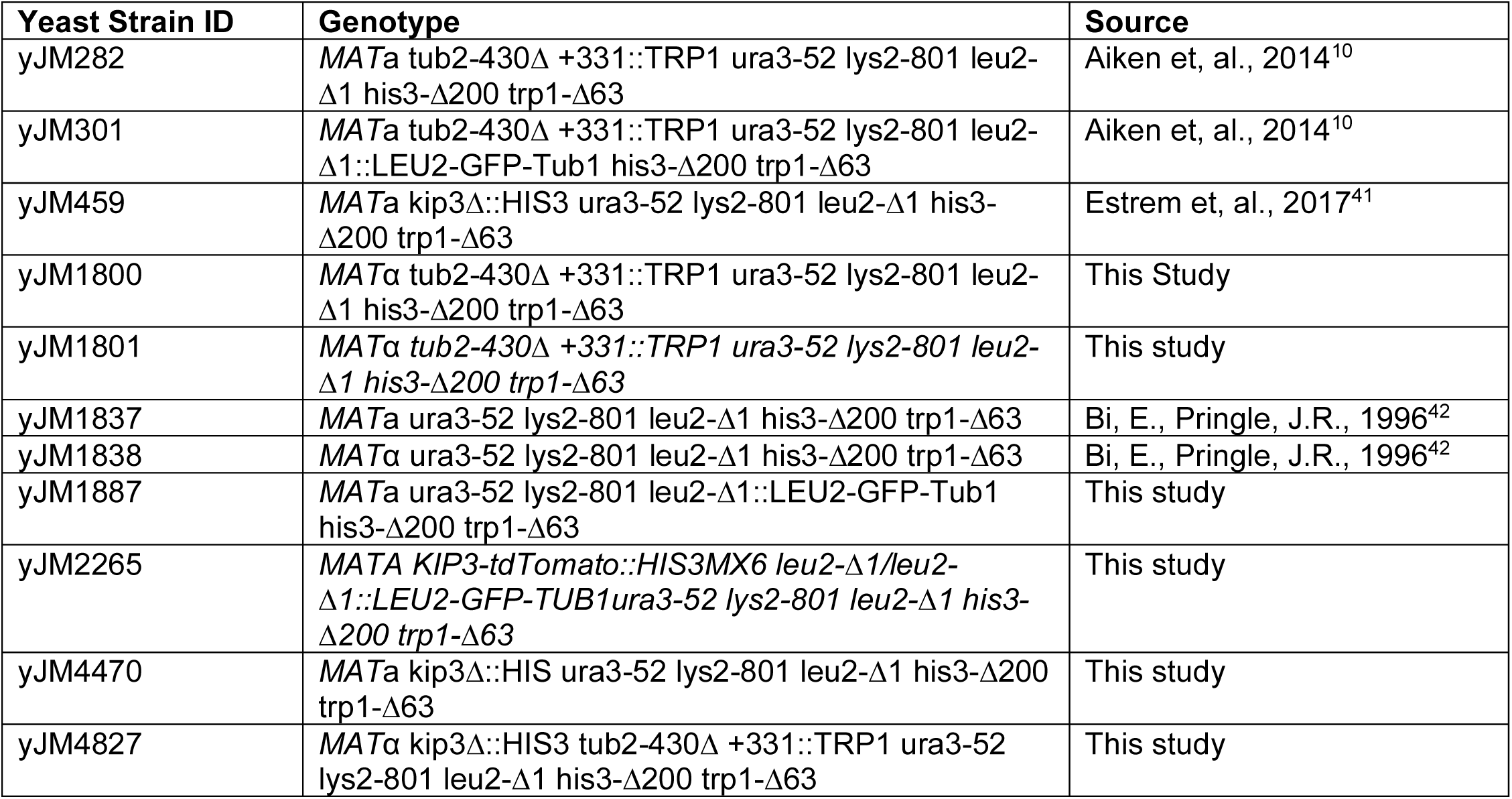
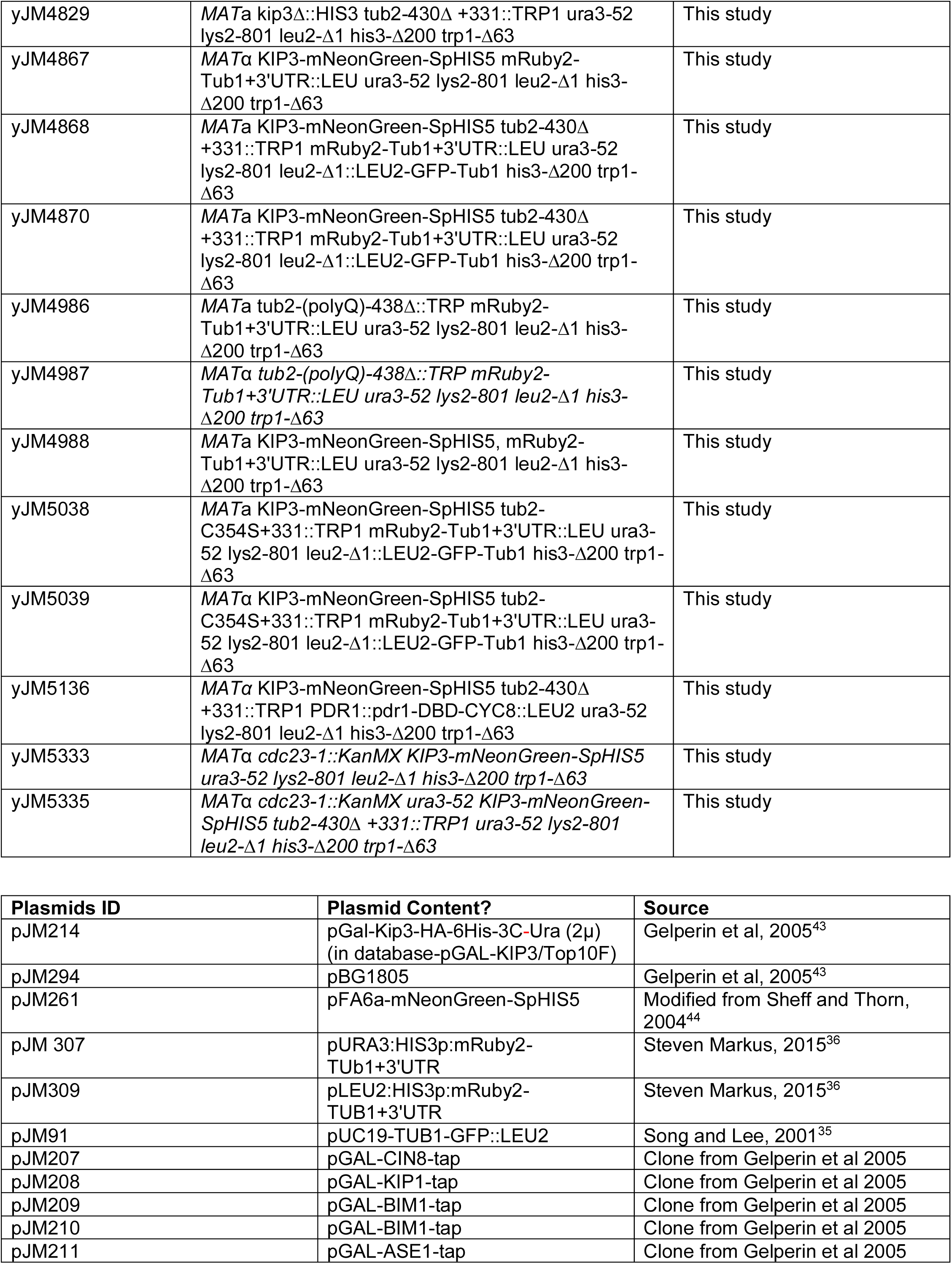
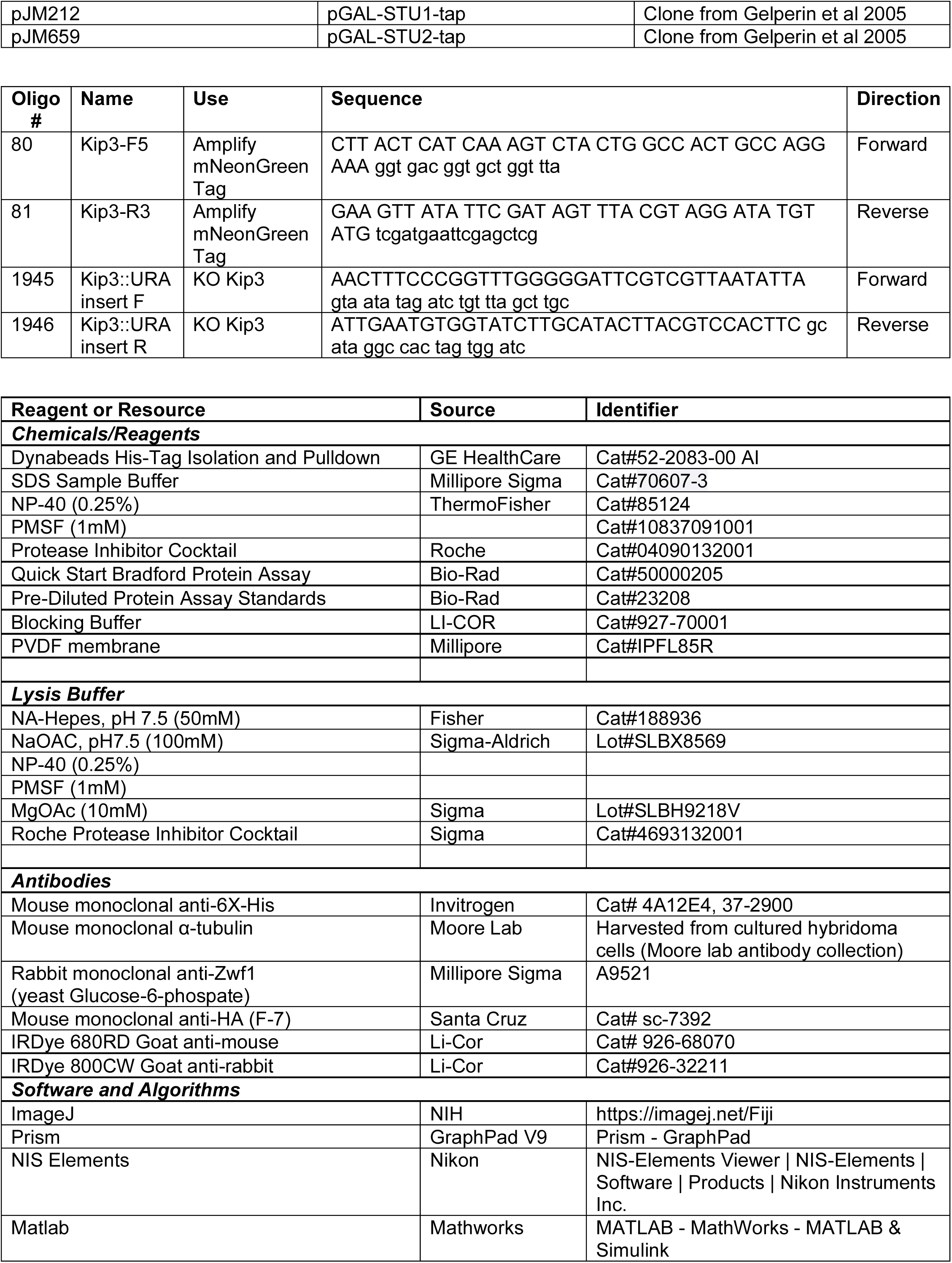

**Supplemental 1: β-CTT is required for Kip3 function**

**(A-B)** The 22 overlapping genes were further sorted by gene ontology using the Gprofiler program and the top 5 biological processes and cellular components are listed. Numbers listed next to bars depict the number of genes associated with the function. **(C)** Example images of wild-type, *kip*3Δ, *tub*2-430Δ, and a *kip*3Δ/*tub*2-430Δ double mutant strains tagged with GFP-Tub1 and imaged during anaphase via confocal microscopy at 30°C. **(D)** Quantification of spindle lengths measured from each end of the spindle pole bodies. A one-way Anova with a Tukey’s Post-Hoc test was completed where p<0.05 is significant (*p=0.0317, ****p<0.0001, n ≥ 25). Error bars represent the mean with standard deviation. All imaging was completed via confocal microscopy.

**Supplemental 2: Quantifications of Kip3 overexpression**

Wild-type and *tub*2-430Δ with the Kip3 overexpression plasmid were induced for 0-5 hours then collected and lysed or fixed. **(A-B)** Example western blot of Kip3 overexpression time course probing for His signal on Kip3, α-tubulin, and Zwf1 as a control. **(C)** Quantification of Kip3-His signal. **(D)** Quantification of α-tubulin signal. Each dot represents the mean with standard deviation from three separate experiments.

**Supplemental 3: Effects of Kip3 and MAP overexpression on yeast MT network**

**(A)** Quantification of the percent of wild-type and *tub*2-430Δ cells with GFP-tubulin blobs from fixation time course experiments seen in figure 2. **(B)** Kip3 overexpression plasmid in wild-type strains labeled with GFP-tubulin and native Kip3-tdTomato. **(C)** Example images of MAP overexpression in wild-type strains labeled with GFP-tubulin. All imaging was completed via confocal microscopy after 3hrs of induction.

**Supplemental 4: Quantification of Kip3 localization**

Quantifications of Kip3-mNeonGreen in wild-type, *tub*2-430Δ, *tub*2-438Δ, and *tub*2-C354S strains tagged with mRuby-TUB1 (Figure 4). **(A)** Line scan analysis of MT length in microns (****p<0.0001, ***p=0.0005). **(B)** Line scan analysis of Kip3-mNeonGreen intensity across the entire MT lattice (****p<0.0001). **(C-D)** Kip3-mNeonGreen intensity was measured by drawing 7×7 boxes at the **(C)** plus end of the MT (**p=0.0013) and **(D)** the middle of the MT (****p<0.0001) as determined by the mRuby-tubulin signal. A one-way Anova with a Tukey’s Post-Hoc test was completed for all graphs where p<0.05 is significant. Bars represent the median value.

**Supplemental 5: Quantification of Kip3 and tubulin pull-down**

The Kip3 overexpression plasmid was transformed into wild-type and *tub*2-430Δ strains and ectopic Kip3 was induced for 2 hrs. **(A)** Quantification of Kip3 intensity (p=0.3274). **(B)** Quantification of tubulin binding. Empty plasmid control was included in experiments to measure tubulin binding to the beads. A one-way Anova with a Tukey’s Post-Hoc test was completed where p<0.05 is significant (*p=0.0175, **p=0.0091).

