## Supplementary figures and images for "Kinesin-8/Kip3 requires beta tubulin tail for depolymerase activity"

### Supplemental Figure 1

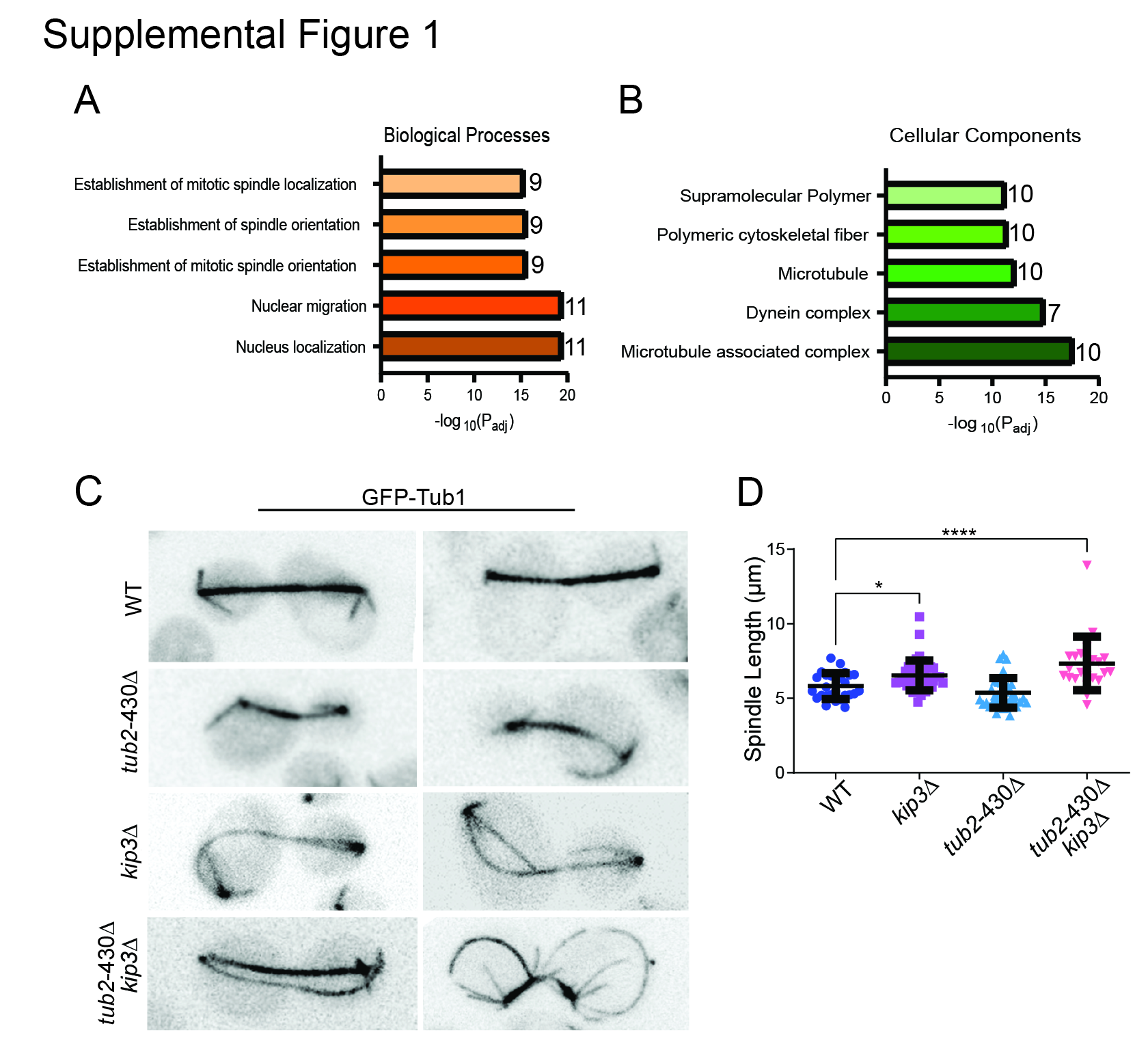

### Supplemental Figure 2

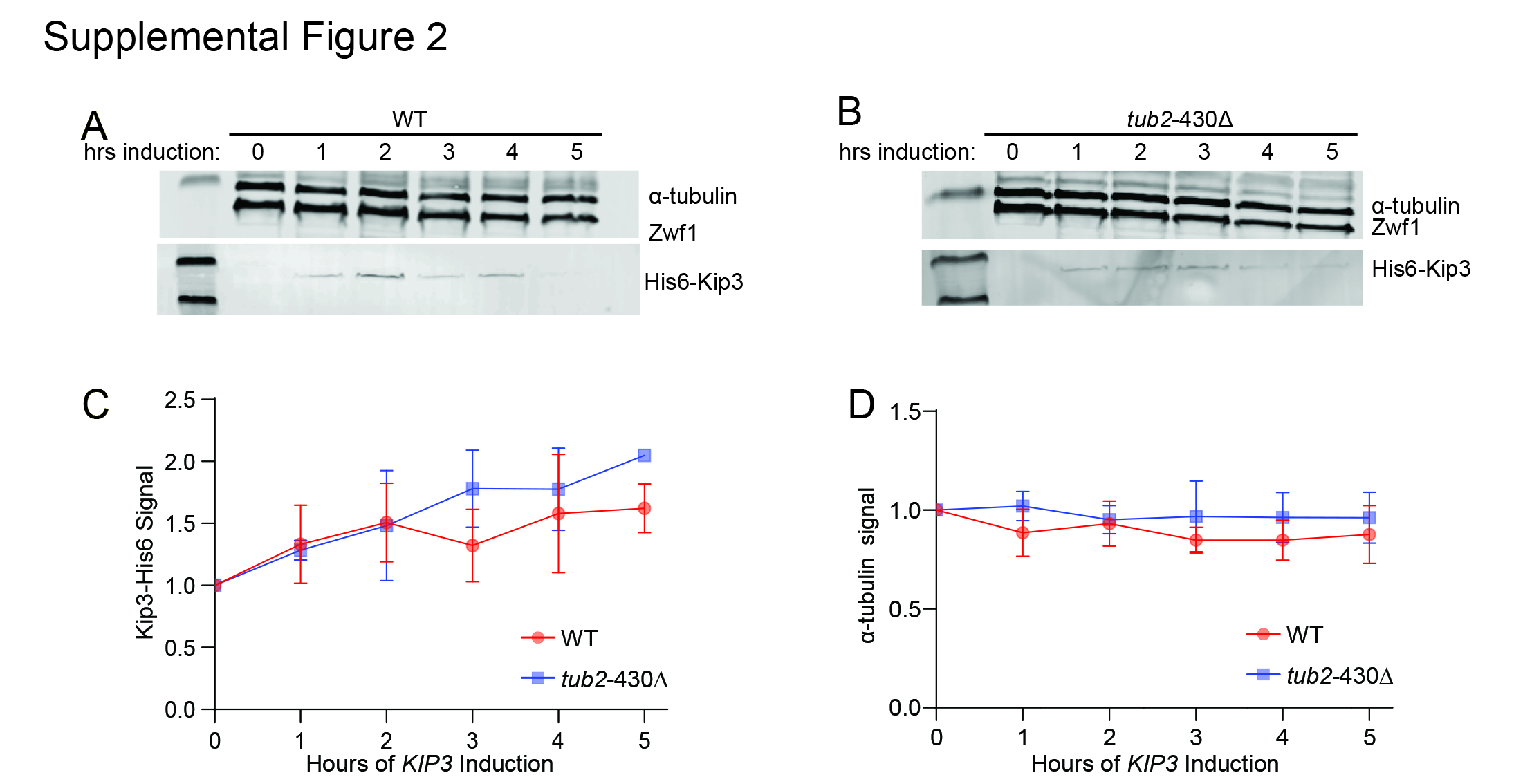

### Supplemental Figure 3

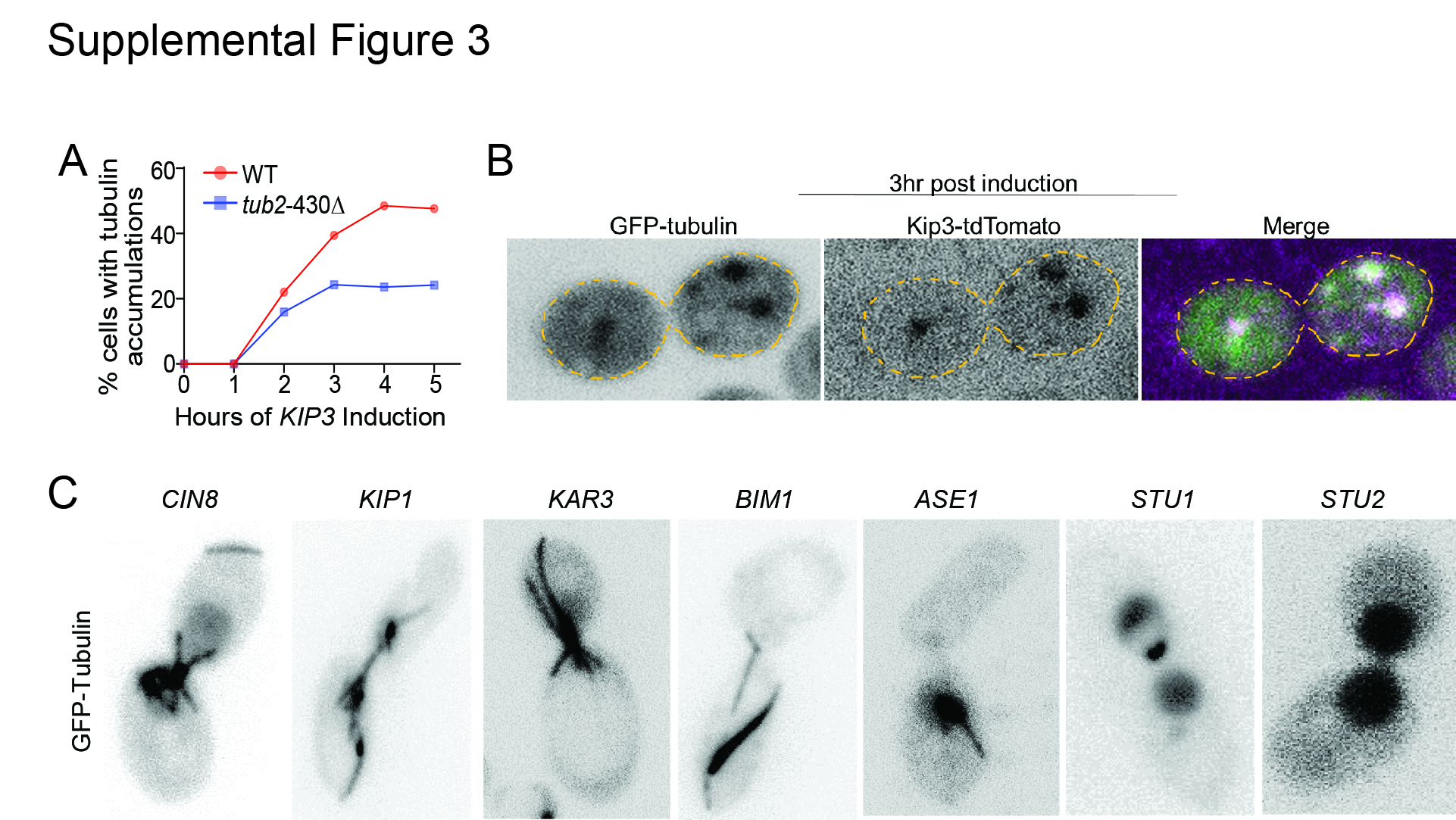

### Supplemental Figure 4

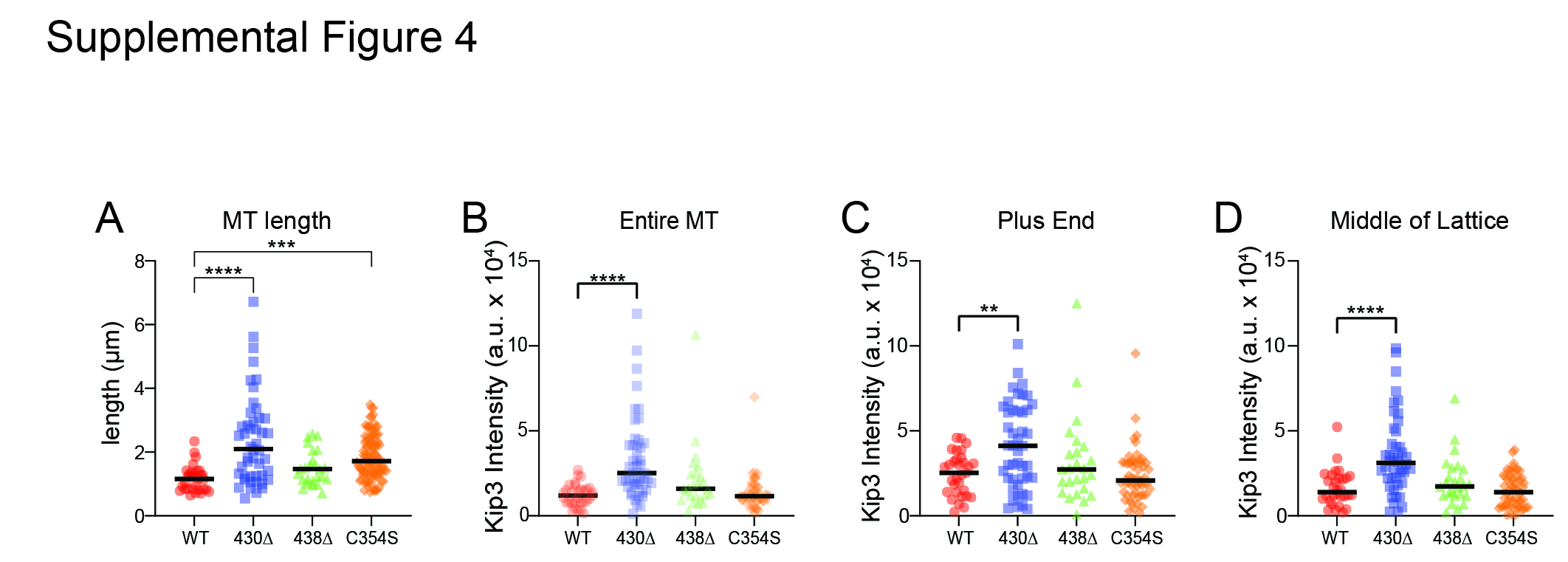

### Supplemental Figure 5

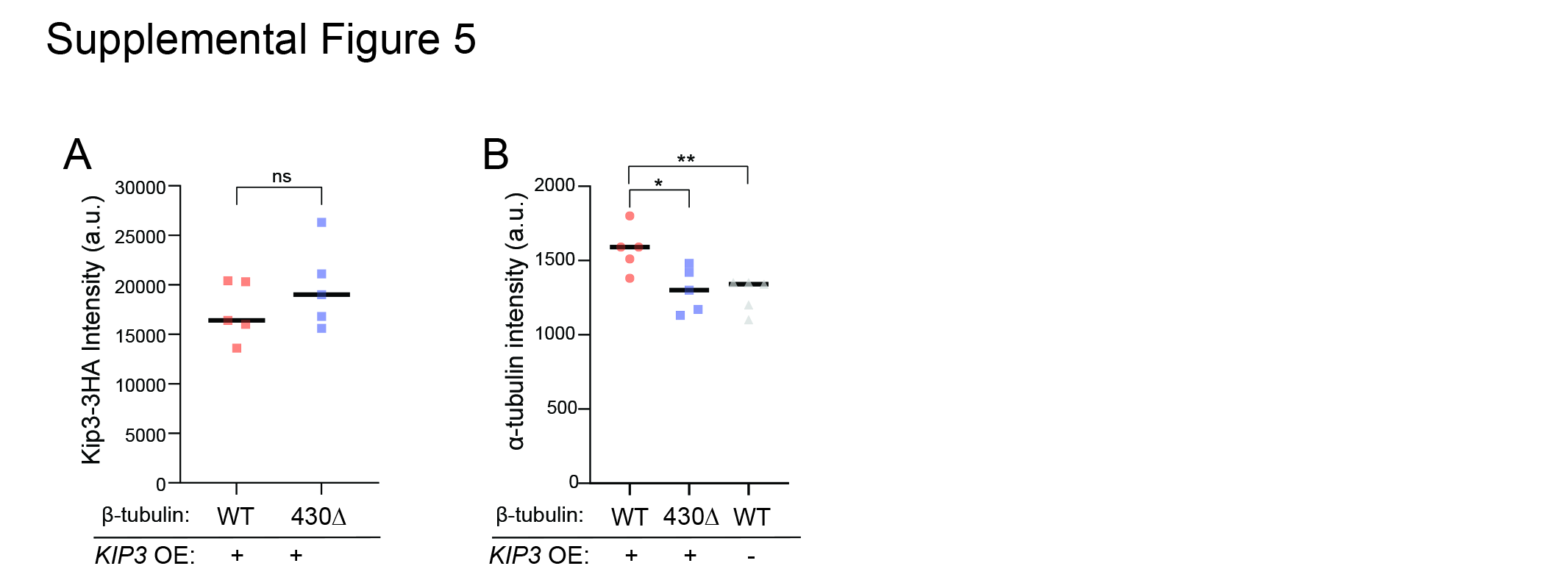
